# Deconstructing *taxa x taxa x environment* interactions in the microbiota: A theoretical examination

**DOI:** 10.1101/647156

**Authors:** Senay Yitbarek, John Guittar, Sarah A. Knutie, C. Brandon Ogbunugafor

## Abstract

1. A major objective of microbial ecology is to identify how the composition of gut microbial taxa shapes host phenotypes. However, most studies focus solely on community-level patterns and pairwise interactions and ignore the potentially significant effects of higher-order interactions involving three or more component taxa.
2. Studies on higher-order interactions among microbial taxa are scarce for many reasons, including experimental intractability, daunting diversity and complexity of many microbial systems, and the potential confounding role of the environment. Moreover, we still lack the empirical and statistical tools to isolate and understand the role of higher-order interactions on the host.
3. Here, we apply a mathematical approach to quantifying the effects of higher-order interactions among taxa on host infection risk. To do so, we adapt the Hadamard-Walsh method recently used in evolutionary genetics to quantify the nonlinear effects of mutations on fitness. We apply our approach to an *in silico* dataset built to resemble a population of insect hosts with gut-associated microbial communities at risk of infection from an intestinal parasite. Critically, we examine these interactions across a breadth of environmental contexts, using nutrient content of the insect diet as a model for context.
4. We find that the effect of higher-order interactions is considerable and can change appreciably across environmental contexts. Strikingly, the relative eminence of different orders (pairwise vs. third order, fourth order, and fifth order) changes as a function of environmental context. Furthermore, we show–in our theoretical microcosm–that higher-order interactions can stabilize community structure thereby reducing host susceptibility to parasite invasion.
5. Our approach illustrates how incorporating the effects of higher-order interactions among gut microbiota across environments can be essential for understanding their effects on host phenotypes. We conclude that higher-order interactions among taxa can profoundly shape important organismal phenotypes, and they deserve greater attention in host-associated microbiome studies.

## Introduction

Animal guts contain complex microbial communities whose structure and function depend upon the interactions among microbes and the host. Gut microbiota serves as key actors in host health, impacting development, metabolism, and the immune system (Brugman et al., 2018; McFall-Ngai et al., 2013). The development of axenic and gnotobiotic model hosts has made it possible to experimentally study how the microbiota influences host traits of interest (Douglas, 2018). However, most studies rely on correlations between the relative abundances of individual microbial taxa and host traits (e.g. immune function), and also community-level patterns at family level taxonomic resolutions, ignoring the potential influence of higher-order interactions among taxa within the community (Hooper et al., 2012; Knutie et al., 2017; Macpherson & Harris, 2004; Round & Mazmanian, 2009).

The field of complex systems is increasingly interested in understanding the emergent properties of higher-order interactions (Battiston et al., 2020). Higher order interactions have been the object of relatively rigorous inquiry in the realm of genetics, where they are discussed in terms of epistasis, or non-linear interactions between genes and mutations (Mackay & Moore, 2014; Weinreich et al., 2013, 2018a). A useful non-technical definition of epistasis is “surprise at the phenotype when mutations are combined, given the constituent mutations’ individual effects” (Weinreich et al., 2013). In particular, higher-order epistasis is of interest, as these interactions comprise all of the complexity and challenges of understanding and studying higher-order interactions in other systems, and even in microbes (Gould et al., 2018). Not unlike genomes, communities or neural circuits, insect gut microbiomes are complex systems defined by the interaction between individual entities or parcels of information (in this case, component taxa in the microbiota). Consequently, we might predict that higher-order interactions between taxa in the microbiota might underlie microbiota-associated organismal phenotypes.

A long-standing goal of ecology is to capture the vast diversity of multispecies interactions—the unpredictable effects that arise when multiple species are present in an ecosystem (Barabás et al., 2016; Chesson, 2000; Hutchinson, 1961; Mayfield & Stouffer, 2017; Vandermeer, 1969). For example, animals harbor diverse microbial communities that are variable in their composition, governed by stochastic processes, which influences the overall behavior of the system (Douglas, 2018). This problem has more recently become the object of inquiry in communities of microbes (Enke et al., 2019; Guittar et al., 2019; Mickalide & Kuehn, 2019; Sanchez-Gorostiaga et al., 2018). Many ecological studies involving complex network structures typically focus on pairwise interactions (Kareiva, 1994; Levine et al., 2017; Mayfield & Stouffer, 2017). Only very recently has the literature demonstrated that higher-order interactions are at play in these systems, an important area for further inquiry, given how they may potentially complicate (or even undermine) simple models of microbial community function (Sanchez-Gorostiaga et al., 2018).

Higher-order interactions in the gut microbiota of *Drosophila specie*s impact lifespan, fecundity, development time, and community composition (Gould et al., 2018). With a gut community comprising five core taxa, Gould et al. found that three-way, four-way, and five-way interactions accounted for 13-44% of all possible cases depending on the host trait. Yet, lower-order interactions (2-pairs) still accounted for at least half of all the observed phenotypes in the system. Work by Sanchez-Gorostiage et al. (2018), examined the contributions of multispecies interactions to determining community function (i.e. amylase expression). In the presence of higher-ordered interactions, the predictive power of the additive null model (absence of interactions) in predicting community function decreases. However, by accounting for both behavioral and population dynamics effects into their null model, higher-order interactions did provide good predictions for community function. Higher-order interactions can have important implications for the predictive power of bottom-up approaches to designing complex communities and determining their functional traits (Sanchez-Gorostiaga et al., 2018). The aforementioned studies provide examples of how higher-order interactions can be measured and suggest that they are relevant for understanding how microbial taxa influence certain phenotypes. While the importance of diversity and host interactions is clear, to our knowledge no studies have attempted to specifically disentangle effects of higher-order interactions across environmental contexts.

One major barrier to more of these studies is the paucity (or non-existence) of the datasets structured like those in an evolutionary genetics framework, such that existing statistical methods might be used to resolve interactions (Tekin et al., 2017; Wood et al., 2012). For example, the problem of constructing a set of insects that each carry a different combination of constituent taxa of interest grows exponentially with the number of taxa. And unlike some genetic systems, constructing a different insect with a different set of bacterial taxa (corresponding with the possible combinations of taxa) is currently a non-trivial technical challenge. Nonetheless, the use of combinatorial complete datasets—insects containing all combinations of taxa (even few in number)— to explore higher-order interactions (beyond a single taxon or pairwise interactions) could help to inform how taxa interact in framing organismal phenotypes. Higher-order interactions could, in principle, be used to examine how our predictions for taxa-taxa interactions will be contingent on the host context in which a certain distribution of taxa exists.

In this study, we reframe how we consider higher-order interactions in an insect gut using theoretical approaches. We apply a relatively simple mathematical method called the Walsh-Hadamard transform (WHT), which has been used to demonstrate how higher-order interactions between mutations influence fitness or other organismal traits (Poelwijk et al., 2016; Weinreich et al., 2013, 2018b). We use this method to explore how higher-order interactions among gut taxa can influence host fitness, across micro-environments. In this study, we use it to quantify higher-order interactions in an *in silico* dataset resembling the type of data that can be collected, presently in genetic systems, and plausibly in the future in microbiota experimental systems.

We have chosen to consider the nutritional environment of the host, as resources can vary due to spatial and temporal differences, and in terms of the quantity and quality of required resources. A key component of resource availability is nutrition, which is likely to influence host resistance to natural enemies. In microbial systems, increased resource availability resulted in greater host resistance to parasites (Gómez et al., 2015; Lopez-Pascua & Buckling, 2008). Lower resource levels have been found to be costly for resistance to parasites in Drosophila melanogaster (McKean et al., 2008). Nutritional content (quality and quantity) is a well-known stressor for insect microbes in many settings, including the gut microbiota (Engel & Moran, 2013; Gurung et al., 2019; Mereghetti et al., 2017; Skidmore & Hansen, 2017). However, experimental studies involving model systems rely on high nutritional diets to understand factors affecting susceptibility to infectious diseases (Roberts et al., 2019). In this work, we consider how varying nutritional environments influence host susceptibility to disease risk.

Using this framework, we are able to examine underappreciated aspects of the microbiota: questions surrounding the notion that the microenvironment of the insect gut may shape higher-order interactions between taxa, with important consequences for host health and fitness. Our study examines the consequences of higher-order microbial interactions for host susceptibility (i.e. phenotype of interest) to disease risk. We hypothesize that higher-order interactions underlie host microbiome robustness to intestinal parasite invasion, reducing host susceptibility to disease risk, and that these interactions are highly dependent on environmental context. While this study is designed to address standing questions about interactions within the microbiota, it also offers future directions. We introduce this approach with the hope that it, or a related method may eventually be applied to a tractable experimental system for real-world validation and believe that insect systems are among the most promising candidates for these examinations.

## Methods

### Data source

The data used in this study arise from raw data used to generate theoretical fitness landscapes, composed of five-bit strings that were generated from an *in silico* data set introduced in a prior study (Meszaros et al., 2019). The data set was originally generated in order to provide large empirical data sets that could be used to study advanced topics in population genetics, including higher-order epistasis. The datasets are constructed such that they can serve as an exploratory space for any theoretical set of interactors, and therefore, is well-structured for the study of interacting microbial taxa. That is, there is nothing about the datasets that renders it a better fit for any one biological problem than another: these data could just as well be used to study interacting genes as taxa or any parcel of information. The data are defined as strings of information (e.g. 01011 or 11001), each with a corresponding “phenotype” value. Therefore, this data is equipped for the analyses as proposed in this study. Here, we use it to generate theoretical microbiota in an insect gut. For more information on the data set and its origin, see Meszaros et. al 2019 and the Supplementary Information.

For the purpose of this study, it is important that we are transparent with regards to the data source, the notation, and the method for transforming the data into a microcosm for taxa in an *in silico* insect microbiota. In this study, our hypothetical insect guts are encoded as strings of bits. Bits can either be 0 or 1. 1 indicates the presence (+) of a taxa. 0 indicates absence (-) of a taxon. For example, we can write a string of 0 and 1 corresponding to an insect gut with five interacting taxa (A-D) as demonstrated in Table 1.

**Table 1.**
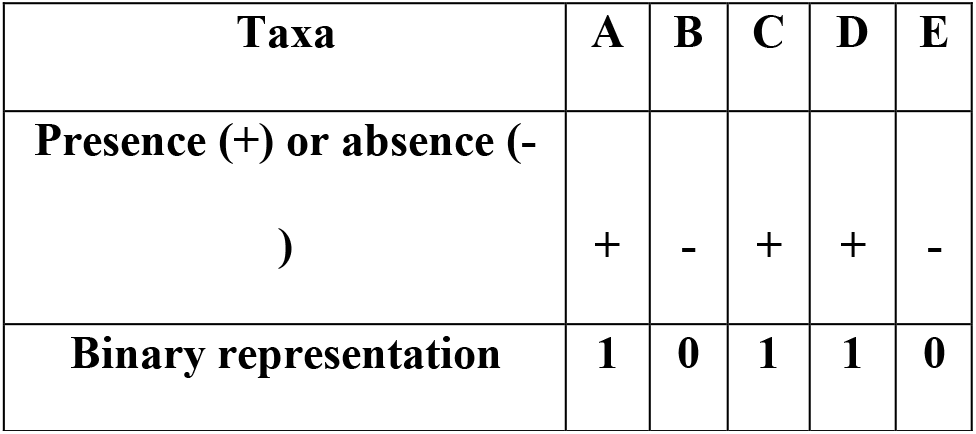
Data structure of a hypothetical insect gut.

As we can see, the “10110” string corresponds to an insect gut where taxa A is present, B is absent, C is present, D is present, and E is absent. The data set that we are mining, originally derived for studying combinatorial data sets that are common in the study of fitness landscapes, offers tens of thousands of combinatorial sets that correspond to a hypothetical insect gut with interacting taxa (Meszaros et al., 2019). We have randomly chosen one such set, containing five individual bits, to explore the central biological concepts of interest in this study: the measurement of higher-order interactions between taxa, and how these interactions might be influenced by the environmental context.

The data set that we will use is a five-bit string (a string of five numbers), combinations of presence and absence (+ or -) of five taxa (A-D). The combinatorial possibility corresponds to 2^5^ = 32 theoretical combinations of taxa across four different insect environments. In Table 1, we show the “fitness” (host infection risk in our study) values for all 32 *in silico* combinations of taxa across four different insects.

### Calculating the strength of interactions

As mentioned in the introduction, there are myriad methods for resolving higher-order interactions, and many such methods have been explored in genomic studies (Crona, 2020; Domingo et al., 2018; Guerrero et al., 2019; Otwinowski et al., 2018; Poelwijk et al., 2016; Sailer & Harms, 2017). A full treatment of the strengths and weaknesses of every method would require a review that is beyond the scope of our study, but some existing work has interrogated multiple methods in the study of epistasis (Poelwijk et al., 2016). In describing the methods as applied in this study, we have erred on the side of redundancy in our explanations. We believe that this is appropriate, given that our method of choice – the Walsh Hadamard Transform – has never been applied to the study of the microbiota and so could benefit from further explanations.

### The Walsh-Hadamard Transform

The Walsh-Hadamard Transform allows one to quantify the eminence of interaction effects among potentially interacting objects or parcels. Its main output is a Walsh coefficient, which communicates the magnitude (how large the interaction is) and sign (positive interaction or negative direction) for a given interaction. The method implements phenotypic (host infection risk in our study) values in the form of a vector, before reformatting it into a Hadamard matrix (and is then scaled by a diagonal matrix). The output is a collection of coefficients which correspond to the strength of interaction between taxa.

For example, we can define the Walsh Hadamard coefficient for the following:

<u>*</u>B<u>*</u>DE

The asterisks (<u>*</u>) correspond to taxa that could either be present or absent. This can reencoded in binary as:

01011

This Walsh Hadamard coefficient for this string would correspond to the magnitude of the interaction between the B, D and E taxa. Importantly, we would label the interaction between B, D and E as a “third order” interaction, as the calculation provides the average strength of interaction between three different taxa: B, D and E. Understanding the different orders of interaction is the key to gaining a perspective on “higher-order” interactions. In a gut containing five taxa that we are interested in understanding the interaction between, there are five different “orders” of potential interaction.

For example:

0^th^ (zeroth) order interaction would be the insect containing none of the taxa of interest (A-E) present. First order interactions correspond to the influence of individual taxa on the infection risk. There are five such first order terms in this theoretical insect microbiota:

A****

*B***

**C**

***D*

****E

Similarly, there are ten second order coefficients, ten third order, five fourth order, and one fifth order (corresponding to the interaction between all five taxa; ABCD or 11111). These Walsh Hadamard coefficients can be summed within an order. Consequently, a whole theoretical “insect gut” can be described in terms of the overall magnitude of its 0 – 5^th^ order interactions. For example, we can examine the strength of third-order interactions (in sum) and compare them to the strength of fourth order interactions.

The Walsh Hadamard coefficient describes the magnitude to which an interaction map is linear, or second order, third, and so forth. We refer interested readers to two published manuscripts—Poelwijk et al. (2016) and Weinreich et. al. (2013)—that outline and apply the method in good detail. Also, see the Supplementary Information for a brief primer.

The Walsh-Hadamard Transform relies on the existence of combinatorial data sets, where the objects for which we are interested in understanding the interactions between (taxa in this study) are constructed in all possible combinations. Another limitation of the Walsh-Hadamard Transform is that it can only accommodate two variants per site, that is, two states per actor. In the case of taxa, we can think of this in terms of the presence/absence of a certain taxon, and we can encode this in terms of 0 (absence) or 1 (presence). For each hypothetical insect with a different presence/absence combination, we have a corresponding phenotypic measurement (e.g. host infection risk). For example, if we wanted to measure the higher-order interactions between 4 taxa within an insect with regards to their role in parasite load (as a model phenotype), we would need 2^*L*^ = 16 individual measurements (insects in this case), with *L* corresponding to the number of different taxa whose effects we were interested in disentangling. We can encode this combination of 4 taxa in bit string notation (see Figure 1).

**Figure 1.**
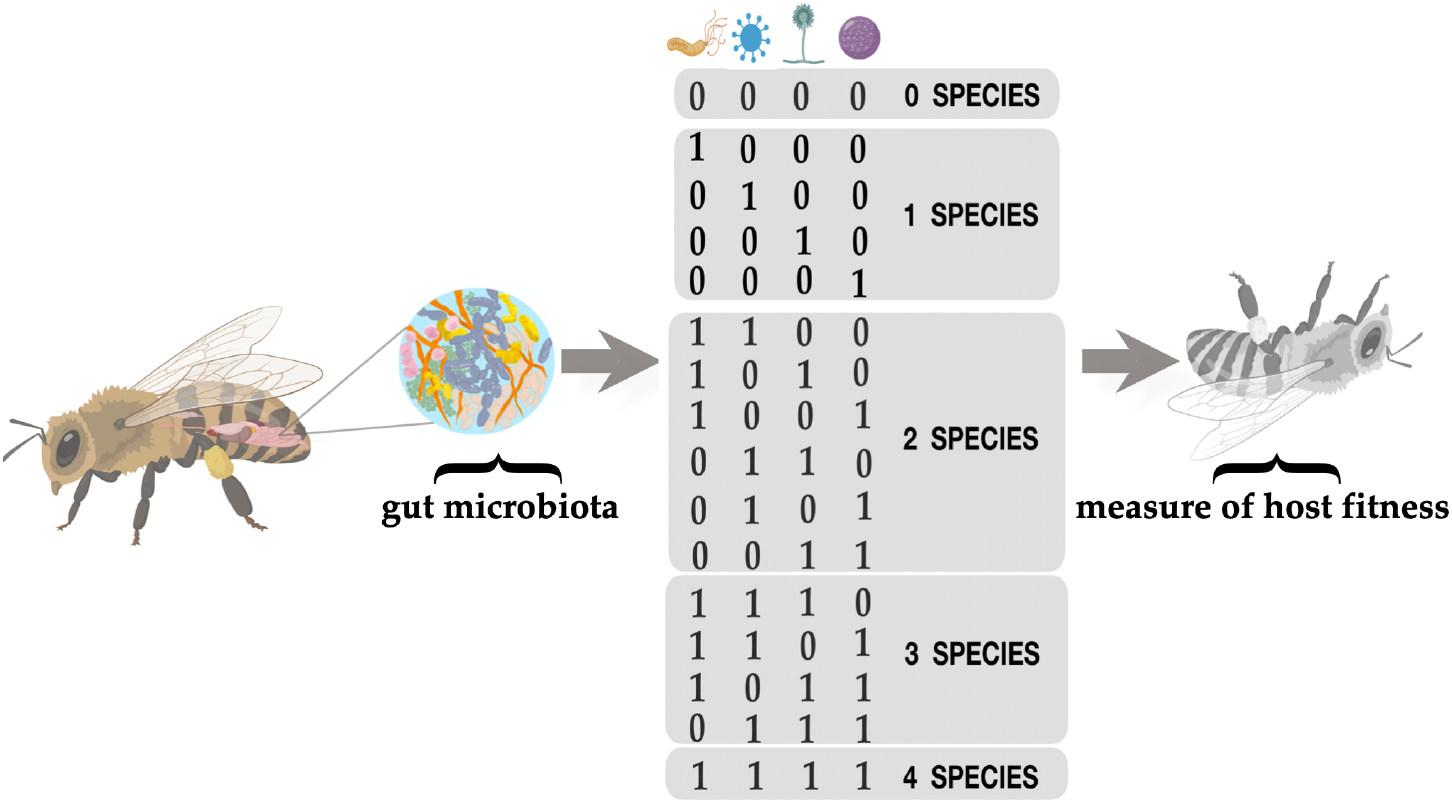
Schematic representation of higher-order interactions in the insect gut microbiota. We represent the presence of microbial species in the gut similarly to the presence of a genetic locus. Species composition of gut microbiota is represented in binary strings. In this configuration, the combination 0011 represents both the presence and absence of two species. For each string combination, we associated a phenotypic measurement, such as infection risk. We quantify “epistatic” interactions between microbes in *n* dimensional space, where *n* represents the number of species interacting.

As described above (Methods section), each site (0 or 1) in the string corresponds to the presence or absence of a given taxa in a given insect. This notation allows us to keep a mental picture of which taxa are in which insect for which we have a phenotypic measurement and can be used to construct a vector of values. Again, the string 01011 corresponds to an insect with the pattern of absent (0), present (1), absent (0), present (1), present (1). The full data set includes a vector of phenotypic values for all possible combinations of taxa—(see Table 1). Note, again, that these can be divided into different classes based on the “order” of the interaction. This vector of phenotypic values for the 32 will be multiplied by a (32 x 32) square matrix, which is the product of a diagonal matrix *V* and a Hadamard matrix *H*. These matrices are defined recursively by:

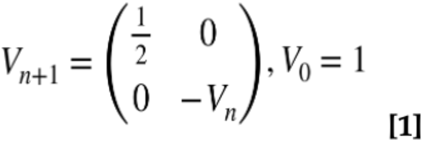

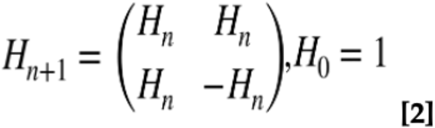

*n* is the number of loci (n = 4 in this hypothetical example). This matrix multiplication gives an output:

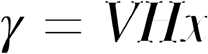

Where *V* and *H* are the matrices described in [1] and [2] above, and γ is the Walsh coefficient, the measure of the interaction between parcels of information in a string. Using this, we compute γ values for every possible interaction between bits in a given string. These methods measure every one of these interactions (e.g. all ten 2nd order interactions) between taxa. While our use of a five-bit string structure (as opposed to an three or fifteen bit string) is arbitrary, it communicates the nature of the higher-interaction problem: Even if we suspect that only five taxa in an insect microbiota are meaningfully influencing a phenotype of interest (Cagnolo et al., 2011; Ferrari & Vavre, 2011; McLean et al., 2016), the possible ways that these species are interacting, and the number of measurable coefficients between them can be meaningful.

Having outlined the method used to quantify higher-order interactions above, it is important to directly explain the presumptive biological interpretation of the values. The Walsh Hadamard Transform returns a Walsh coefficient for each “order” of interaction. This corresponds to the relative strength or importance of that “order” in the phenotype being measured. Therefore, the Walsh-Hadamard Transform can help to interpret the overall presence and eminence of higher-order interactions between taxa in a microbiota.

### The theoretical environment of the insect gut microbiota

Here, we explore how varying nutrient diets influence host susceptibility to parasites in the gut microbiota. We chose to focus on the nutrient diet content in our study design because the resource environment is highly relevant to the insect gut microbiota. In insects, nutrition content of the host’s food can be controlled by the addition of methyl cellulose (an indigestible bulk agent) in the standard food medium (Boots & Begon, 1994). Resource-levels varying from high-quality diets (containing no methyl cellulose in the food medium) to lower-quality diets (replacing 10%, 20%, 30%, 40%, 50%, 60%, 70%, 80%, 90% of the food medium with methyl cellulose) have been utilized to empirically study the role of varying nutrition environments to parasite resistance in lepidopteran pest species (Boots et al., 2011). In our theoretical study, we define “nutrient content” as a diet compromising a range of nutrients in a standard insect diet. A diet of 0 % would correspond to an extremely low nutrition diet, and 100% to a high-quality diet composed of the standard food amount for insects. Consequently, the nutrient gradient 0 – 100% represents varying degrees of resource availability.

## Results

### Norm of reaction

The norm of reaction demonstrates that two insect guts, corresponding to 00000 (no taxa) and 11111 (the presence of taxa of every kind) have the largest parasite loads relative to other insect microbiota combinations. The high parasite load pattern is consistent across the nutrient content that insects consume (Fig. 2). In contrast, we find that parasite load is drastically reduced for all other insect microbiota combinations (examples include combinations 001100; 11011; 11101).

**Figure 2.**
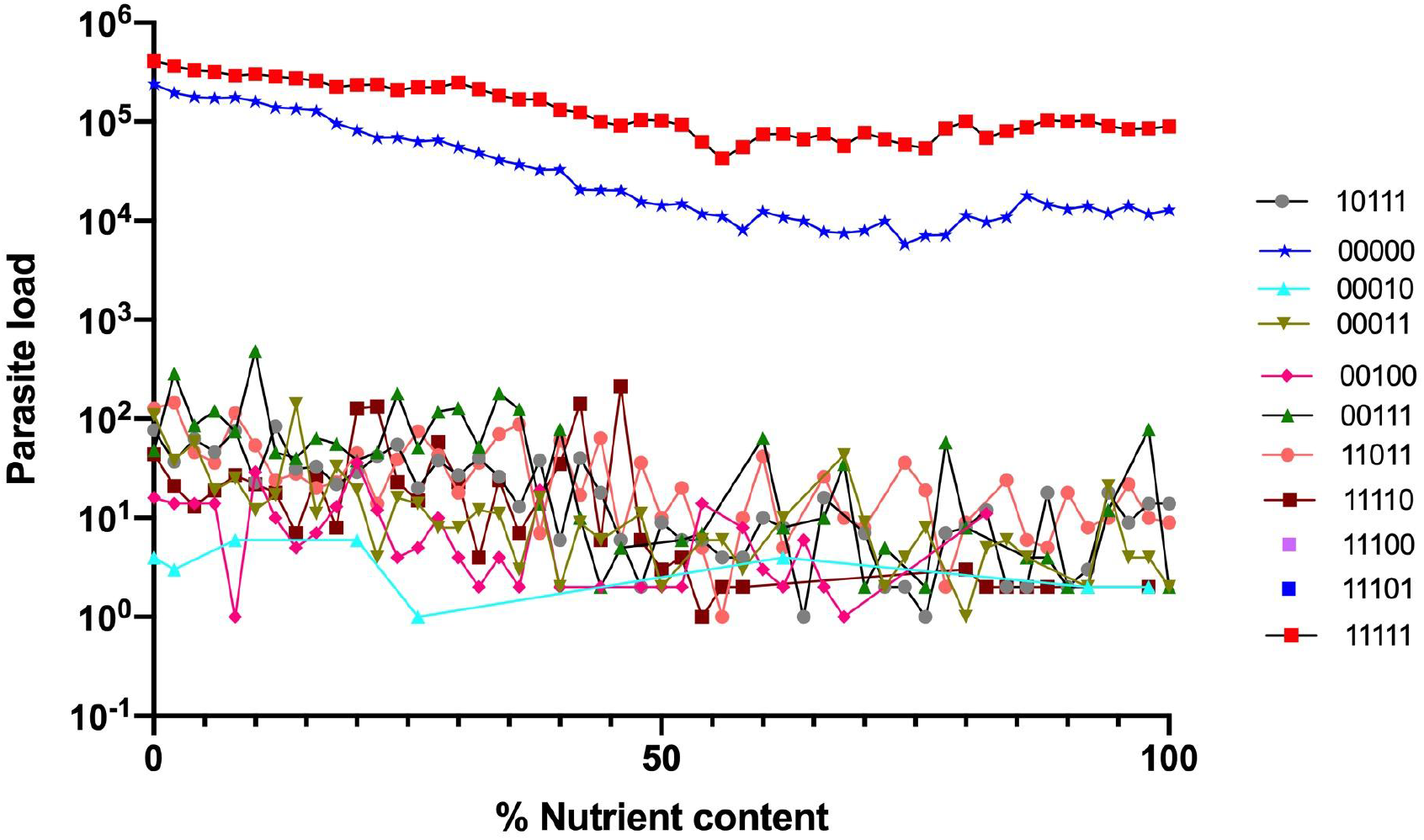
A Norm of reaction representing the parasite load of in silico insect guts as a function of nutrient microenvironment. The *x-axis* represents the nutrient content that insects consume, ranging from 0% (deprived) to 100% (a full, standard nutrient content. Individual data points correspond to insect cuts containing different combinations of taxa. The *y-axis* represents the parasite load, a proxy for the susceptibility of a given insect to infection by parasites. Note that only a subset of the 32 taxa are represented in this, as many of the *in silico* insect guts have parasite loads that are very low. The data for all 32 can be found in the supplementary material.

### Comparison of the orders of interactions among taxa across microenvironments

Figure 3 demonstrates the sum of the absolute values of the interaction coefficients. Here, we can observe the raw magnitude (whether positive or negative in sign) of higher-order interactions as a function of interaction order. Note how the eminence of the higher-order effects changes as a function of nutrient content. At low nutrient contents, fourth order effects are the most impactful on the overall parasite load. At approximately 20%, the fifth order effects (corresponding to the five-way interaction of taxa in the *in silico* insect gut represented by 11111). The change in order of eminence also applies to the second order (pairwise) and third order interactions. At low nutrient contents, the pairwise interactions exert a more meaningful influence on the parasite load than the three-way interactions. At approximately 20% nutrient content--not far from that nutrient percentage where a switch between fourth and fifth order effects manifests--the three-way interactions supplant the pairwise effects in their overall influence on parasite load. Note that all of these values—the *in silico* parasite load data, the interaction coefficients for all individual interactions, and the scaled, absolute value coefficients—can be found in the Supplementary Material.

**Figure 3.**
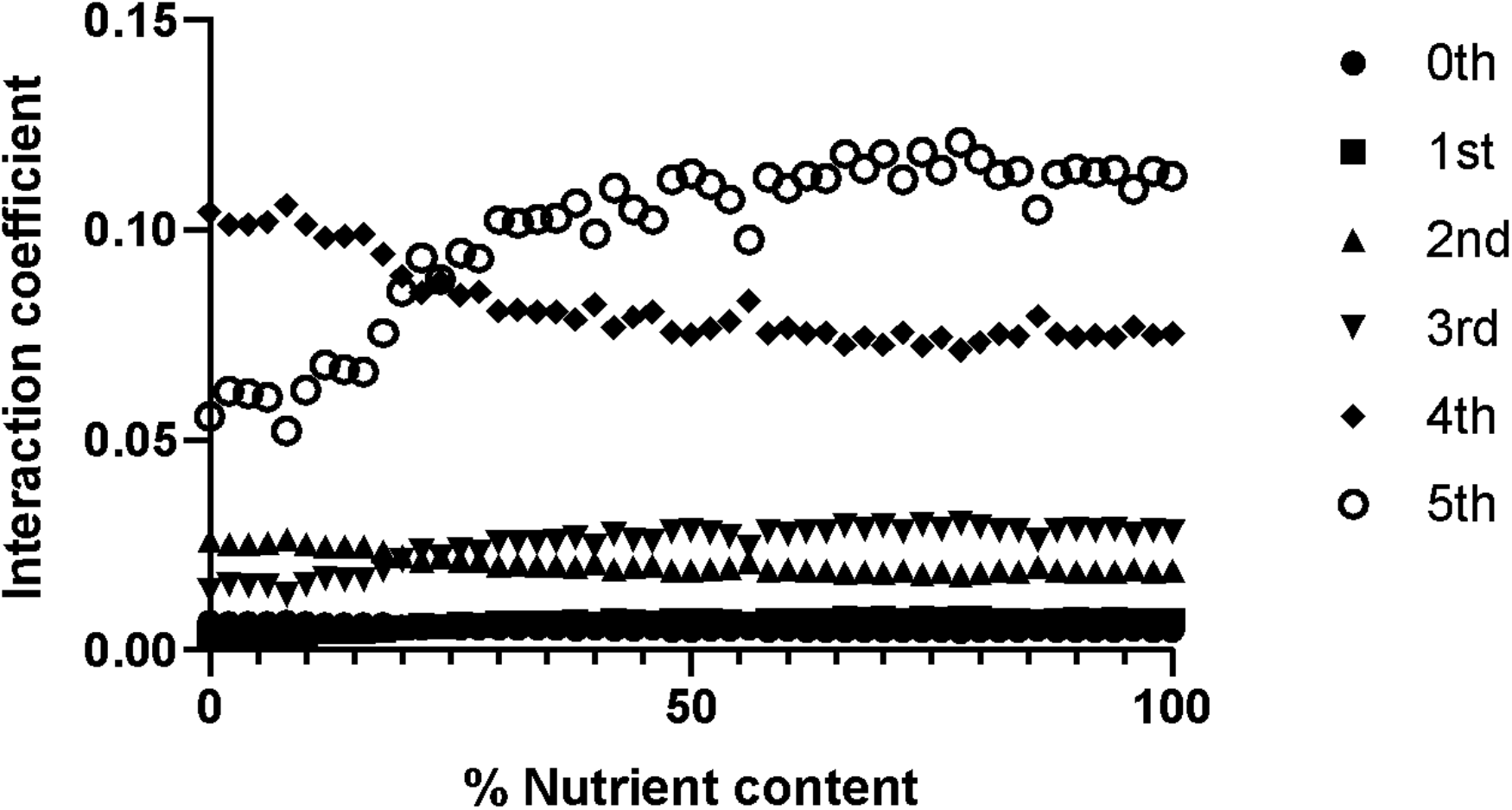
The magnitude of higher-order interaction between taxa as a function of environment (nutrient content). Absolute value, averaged magnitude interactions across interaction orders. The purpose of this depiction is to illustrate how the magnitude (not sign) of the interactions change with interaction order. Of special note is how importance of the order of interactions changes as a function of nutrient content (x-axis). In this scenario, there is a nutrient content threshold (∼20%) where the patterns of the interactions change. In our theoretical insect microbiota system, we define “nutrient content” as a diet compromising a range of nutrients in a standard insect diet. In insect populations, the nutrient content of the host’s food can be controlled by the addition of methyl cellulose (an indigestible bulk agent) in the standard food medium (Boots & Begon, 1994). In our model, a diet of 0 % would correspond to an extremely low nutrition diet, and 100% to a high-quality diet composed of the standard food amount for insects. Consequently, the nutrient gradient 0 – 100% represents varying degrees of nutrient content.

## Discussion

In this study, we explore the possibility of higher-order interactions between taxa that compose an insect gut microbiota. Using *in silico* and mathematical approaches, we demonstrate how higher-order interactions can be measured in a complex system of interacting microbial taxa. In our theoretical scenario, higher-order interactions are present and generally increase in relevance with the order of interaction. Notably, the environment (nutrient content in this case) has a meaningful influence on how higher-order interactions among taxa manifest. This result highlights an aspect of higher-order interactions that is so far largely under-appreciated: that the environment and context in which taxa exist can have a meaningful impact on how taxa interact. Consequently, simply noting that non-linear and higher order interactions between taxa may exist is no longer sufficient in how the insect microbiota is discussed: we must consider, and measure, how environments may influence how interactions manifest. Though our results arise from a theoretical examination of *in silico* insect microbiota, they are results nonetheless (Goldstein, 2018) and highlight the potentially vast scope of the higher-order interaction problem that could define the true dynamics of gut microbiota. Specifically, the outlining of a method that can be used to deconstruct higher-order interactions in biological systems, across environmental contexts, represents a potentially useful contribution to the study of the microbiota.

Though empirical data of the size and scope used in this study are currently challenging to generate, this intractability may be temporary, and future methods may permit the generation of data similar in structure to those explored in our theoretical examination. Note that the calculations of higher-order interactions, and their dynamic nature, can be considered without knowing the specific mechanism that underlies the nature of these interactions, determining the magnitude of coefficients provides relevant information on the eminence of a given order in the microbiota. One additional benefit of these results is that they can identify those settings (combinations between microbiota and a given microenvironment) that should be the focus on mechanistic study. For example, by identifying the taxa involved in large pairwise interactions, one can then examine the mechanistic basis underlying this pairwise interaction through manipulative experiments.

Our results are consistent from recent findings, where diverse communities are more effective at resisting invasions including E. coli invasion of soil communities (Elsas et al., 2012), plant root bacterial communities (Wei et al., 2015), and experimental invasions in bacterial communities (Lu et al., 2018). Collectively these studies show that outcome of invasions are determined by available resources in the microbiota. Our main result showing that higher-order microbial interactions limits the invasion of parasites across nutrient environments is in agreement with studies that interactions are mediated by underlying resource dynamics. The nutritional status of the gut microbiome plays an important role in the health of hosts. Simple gut microbiotas have been engineered to provide hosts with novel functions, such as the ability to use more complex nutrient sources and to fight against pathogens. Recent work by Sun et al. 2020 shows that in Caenorhabditis elegans, the colonization of cellulolytic bacteria enables *C. elegans* to utilize cellulose, an otherwise indigestible carbon substrate. At the community level, cellulolytic bacteria can also support resident bacterial species with additional functional roles, such as the protection by Lactobacillus in the gut against *Salmonella* infection (Sun et al., n.d.). To test our model, insect gut microbiota could be engineered to explore how higher-order microbial endosymbiont interactions protect against pathogen infection by enhancing the nutritional status of the host.

The mathematical approach used in this study—the Walsh-Hadamard Transform—has been previously used by theoretical population geneticists to measure non-linear interactions between mutations (Weinreich et al., 2013). Several empirical data sets in genetics and genomics have demonstrated that the sign of interaction effects can change readily with the identity of the interacting parcels (Guerrero et al., 2018; Weinreich et al., 2013, 2018a). Given this, we predict that the taxa that compose the gut microbiota might be similarly defined by higher-order interactions, and that these interactions will change appreciably with insect microenvironment. The capacity for measuring the effects of higher-order interactions on host fitness is an important step towards understanding the effects of microbiota on their host.

The impact of higher-order interactions in the gut microbiota on host fitness may result from a range of possible interactions, ranging from competitive to mutualistic (Fast et al., 2018; Ludington & Ja, 2020; Newell & Douglas, 2014). To test the full suite of all possible combinatorial interactions and their associated effects on host traits, it is important to experimentally manipulate microbial communities. For example, the fruit fly (*Drosophila melanogaster*) is an attractive model system for designing combinatorial studies due to relative ease of rearing gnotobiotic flies and modularity of its microbiome (Ludington & Ja, 2020). For example, combinatorial designs of microbial communities in *D. melanogaster* revealed that emerging higher-order effects composed of 3, 4, and 5-way interactions impacted aspects of host fitness such as life span and fecundity (Gould et al., 2018). While the relative simplicity and tractability of fly microbiomes facilitates the study of host-microbe interactions, underlying mechanisms can provide insights for more complex mammalian gut microbiomes. In *D. melanogaster*, stable gut colonizers favor specific regions of the foregut, which like mammals, suggest specific niches for gut colonizers (Pais et al., 2018). Therefore, strategies that invertebrates and their microbes employ to form stable associations might be informative for mammalian gut microbiomes (Ludington & Ja, 2020).

## Conclusion

Recent theoretical work suggests that higher-order modeling approaches are able to capture volumes of rich data arising from complex ecological interactions (Battiston et al., 2020). We have adapted approaches from evolutionary genetics to the study of host-associated microbiota. In the future, applying these methods to the analysis of experimental data will yield important insight into microbiome dynamics, towards a richer understanding of just how peculiar the microbiota is, and the many meaningful interactions that it embodies.

## Acknowledgements

We wish to acknowledge the support of organizers and participants of the 2017 RCN-IDEAS arbovirus workshop held in New Orleans. SY acknowledges funding support from NSF Postdoctoral Fellowship award number 1612302. CBO acknowledges funding support from NSF RII Track-2 FEC award number 1736253. The authors would like to thank Victor Meszaros and Miles Miller-Dickson for their input on the *in silico* data, figures and Walsh-Hadamard primer, and Daniel Weinreich for helpful discussion on topics relevant to this study. The authors would like to thank the Associate Editor of J*ournal of Animal Ecology* and two anonymous reviewers for thoughtful feedback on the manuscript. Finally, the authors would like to thank Lawrence Uricchio for helpful feedback on the manuscript.

## Data Availability

The *in silico* data used in this study and code used to generate them can be found on github: https://github.com/OgPlexus/MicrobeTaxa1

## Supplemental Information

The authors can find data, code and other information on: https://github.com/OgPlexus/MicrobeTaxa1.

This also includes a short mathematical primer on the Walsh-Hadamard Transform as applied to binary datasets (also available at https://github.com/OgPlexus/MicrobeTaxa1). For a more rigorous understanding, readers are encouraged to engage the works cited in this manuscript.

## Notes

https://github.com/OgPlexus/MicrobeTaxa1

### Competing Interest Statement

The authors have declared no competing interest.

